# Inter-Organellar Effects of Defective ER-localized Linolenic Acid Formation on Thylakoid Lipid Composition and Xanthophyll-Cycle Pigment De-epoxidation in the Arabidopsis *fad3* mutant

**DOI:** 10.1101/2023.10.09.561516

**Authors:** Monique Matzner, Larissa Launhardt, Olaf Barth, Klaus Humbeck, Reimund Goss, Ingo Heilmann

## Abstract

Monogalactosyldiacylglycerol (MGDG) is the main lipid constituent of thylakoids and a structural component of photosystems and photosynthesis-related proteo-lipid complexes in green tissues. Previously reported changes in MGDG abundance upon stress-treatments are hypothesized to reflect mobilization of MGDG-based polyunsaturated lipid intermediates to maintain extraplastidial membrane integrity. While exchange of lipid intermediates between compartmental membranes is well documented, physiological consequences of mobilizing an essential thylakoid lipid, such as MGDG, for an alternative purpose are not well understood.Arabidopsis seedlings exposed to mild (50 mM) salt-treatment displayed significantly increased abundance of both MGDG and the extraplastidial lipid, phosphatidylcholine (PC). Interestingly, similar increases in MGDG and PC were observed in Arabidopsis *fad3* mutant seedlings defective in ER-localized linolenic acid formation, in which compensatory plastid-to-ER-directed mobilization of linolenic acid-containing intermediates takes place. The postulated (salt) or evident (*fad3*) plastid-ER-exchange of intermediates concurred with altered thylakoid function according to parameters of photosynthetic performance. While salt-treatment of wild type seedlings inhibited photosynthetic parameters in a dose-dependent manner, interestingly the *fad3* mutant did not show overall reduced photosynthetic quantum yield. By contrast, we observed a reduction specifically of non-photochemical quenching (NPQ) under high light, representing only part of observed salt effects. The decreased NPQ in the *fad3* mutant was accompanied by reduced activity of the xanthophyll cycle, leading to a reduced concentration of the NPQ-effective pigment zeaxanthin. The findings suggest that altered ER-located fatty acid unsaturation and ensuing inter-organellar compensation impacts on aspects of thylakoids related to the function of specific enzymes, rather than globally affecting thylakoid function.

**Subject Areas:** (2) Environmental and stress responses

(7) Membrane and transport

## Introduction

Plant responses to salt stress include numerous physiological processes facilitating survival (Park et al. 2016). When exposed to salt, several parallel signalling pathways transduce signals to mediate cytoplasmic or organellar adaptive effects (Park et al. 2016; Wani et al. 2020). As the first layer separating the symplast from the apoplastic environment the plasma membrane has a prominent role in salt and osmotic responses of plants (Amin et al. 2021; Osakabe et al. 2013; Park et al. 2016). Upon salt exposure of the cell surface, cells undergo changes in membrane lipid composition to avoid membrane leakage and maintain cellular integrity (Hou et al. 2016; Munnik and Vermeer 2010; Park et al. 2016). During salt treatment, the plasma membrane experiences a shift in turgor, resulting in bulk-flow endocytosis and large-scale formation of membrane vesicles (Jürgens 2004; König et al. 2008; Mazel et al. 2004). Furthermore, salt induces the formation of ER-plasma membrane contact sites (Lee et al. 2019; Perez-Sancho et al. 2015), which might aid in the redistribution of membrane constituents between organellar membranes to maintain cellular integrity. Previous analyses of salt-treated Arabidopsis plants reported an increased abundance of unsaturated PC upon salt stress (König et al. 2008; König et al. 2007), which likely reflects the processes named above and may be required for the stability of the ER and/or the plasma membrane, either in specialized functional membrane domains or as a large-scale structural membrane element. The molecular details of lipid dynamics conferring salt tolerance to plants at the membrane level are currently not well understood (Wani et al. 2020). While the formation of polyunsaturated fatty acids and their incorporation into membrane lipids has previously been found to contribute to plant salt tolerance (Allakhverdiev et al. 2001; Geilen et al. 2017; He and Ding 2020; Zhang et al. 2012; Zhang et al. 2009), it is largely unclear how membrane lipid unsaturation is remodeled during plant salt acclimation. This study addresses altered formation of PC and possibly other extraplastidial membrane lipids during processes that mobilize lipid intermediates from the plastid.

The plastid is the source of most lipid backbones in plant membranes. The biosynthesis of *cis*-unsaturated fatty acids starts in plastids, where newly synthesized stearic acid is desaturated by the soluble fatty acid desaturase FAB2 to generate oleic acid (18:1^Δ9^; where x:y^Δz^ is a fatty acid containing x carbons and y double bonds in position z counting from the carboxyl end) (Ohlrogge and Browse 1995; Somerville et al. 2000). Oleic acid can be further converted in the plastid by the membrane-intrinsic fatty acid desaturases (FADs) FAD6 (Falcone et al. 1994) and FAD7 (Andreu et al. 2007; Iba et al. 1993) or FAD8 (McConn et al. 1994) to sequentially form linoleic acid (18:2^Δ9,12^) and linolenic acid (18:3^Δ9,12,15^), respectively. A substantial proportion of oleic acid is alternatively exported from the plastid and can be converted to linoleic acid and linolenic acid by the ER-localized (Dyer and Mullen 2001) fatty acid desaturases FAD2 (Okuley et al. 1994) and FAD3 (Browse et al. 1993), respectively. The two compartmentalized pathways for linoleic and linolenic acid formation, one plastidial and one extraplastidial, may have evolved to enable independent control of membrane lipid unsaturation in plastidial and in extraplastidial membranes (Browse et al. 1986; Ohlrogge and Browse 1995; Somerville et al. 2000). Plastidial and extraplastidial lipids differ in their degrees of unsaturation, and a metabolic distinction is in line with reported different roles for polyunsaturated lipids in plastidial photosynthesis (Bastien et al. 2016; McConn and Browse 1998; Vijayan and Browse 2002) and in extraplastidial membrane function (Caiveau et al. 2001; Vaultier et al. 2006). In the present study, we are comparing plastidial and extraplastidial processes by analyzing lipids representative for these compartments, namely plastidial monogalactosyldiacylglycerol (MGDG) and extraplastidial phosphatidylcholine (PC).

MGDG is the main lipid constituent of thylakoids, which are the basis for photoautotrophic sustenance of every green plant. The photosynthetic apparatus within thylakoid membranes consists of protein-pigment complexes using light energy to form NADPH and ATP required for the fixation of CO_2_. In contrast to most other cellular membranes, thylakoids consist mainly of glycoglycerolipids, with MGDG accounting for about 50% and digalactosyldiacylglycerol (DGDG) for 25% of the total lipids in plastidial membranes of green tissues (Boudiere et al. 2014). Compared to the plasma membrane, which displays a mass protein/lipid ratio of approx. 1,thylakoids are more highly saturated with proteins, reaching a mass protein/lipid ratio of up to 1.6. MGDG containing highly polyunsaturated fatty acids is essential for thylakoid function, as photosynthetic complexes require membrane surroundings with particular biophysical properties. Electrons excited by light energy are transported within the photosynthetic complexes, and the electron carriers plastoquinone and plastocyanin enable their movement from one complex to the next. Based on the x-ray crystal structure of photosystem II (PSII) (Umena et al. 2011) and on modeling studies (Van Eerden et al. 2017), it has been proposed that MGDG forms specialized binding sites for protein subunits of PSII that define the orientation of chlorophyll within the complex and contribute to an efficient exchange of electrons between PSII and plastoquinone (Van Eerden et al. 2017). Polyunsaturated MGDG forms inverted hexagonal phases rather than bilayers, enabling rapid lateral diffusion and dynamic protein-lipid interactions important for the photosynthetic apparatus. The precise modulation of thylakoid lipid composition is important for maintaining the integrity of a coherent bilayer of high fluidity to enable the formation of a pH gradient in the thylakoid lumen and, thus, is of key importance for the functionality of the photosynthetic machinery (Kirchhoff 2014; Kobayashi 2016). Inverted hexagonal phases formed by MGDG are also important for the operation of the xanthophyll cycle (XC). It has been proposed that upon illumination a special thylakoid domain is established which consists of MGDG, the light-harvesting complex of PSII (LHCII) and the XC enzyme violaxanthin de-epoxidase (VDE), and that the conversion of violaxanthin (Vx) to antheraxanthin (Ax) and zeaxanthin (Zx) is taking place within this membrane domain (for a recent review, see (Goss and Latowski 2020)).

On the other hand, the glycerophospholipid PC is a main lipid constituent of extraplastidial membranes, such as the ER, Golgi, transitory vesicles, tonoplast or the plasma membrane (Nakamura 2017). PC is the substrate for fatty acid modification in the ER (Somerville et al. 2000) and has previously been implicated in plant stress acclimation (Pical et al. 1999; Tasseva et al. 2004). To maintain the functionality of cellular membranes over the course of development or during acclimation to changing environmental conditions, it is essential for plant cells to be able to modulate the degree of unsaturation in their membrane lipids according to shifting cellular requirements. Changes in membrane lipid composition, possibly reflecting lipid remodeling, are well known for various biotic and abiotic stresses and may be aspects of physiological mechanisms to maintain membrane integrity (Barajas-Lopez et al. 2021; Gigon et al. 2004; Mosblech et al. 2008; Pical et al. 1999). For instance, the degree of membrane lipid unsaturation is essential to maintain the fluidity and integrity of membranes under temperature stress (Wallis and Browse 2002). An increased degree of unsaturation of membrane lipids, such as PC, has also been proposed to mediate salt tolerance (Bonaventure et al. 2003; Feller et al. 2002; Tasseva et al. 2004; van Meer et al. 2008), and in the yeast *Saccharomyces cerevisiae* enhanced membrane lipid unsaturation by itself confers salt tolerance (Zhang et al. 2012). While PC unsaturation changes upon stress according to cellular requirements (König et al. 2008; König et al. 2007), it is currently unclear how this lipid remodeling is achieved. Previous studies have shown that changes in lipid abundance can occur globally within the cells, but so far it is not well studied whether such changes are coordinated between different cellular compartments. Previously reported synchronous changes in the abundance of MGDG and PC in response to salt treatment of Arabidopsis plants (König et al. 2008; König et al. 2007) suggest that lipid intermediates required for plasma membrane remodeling upon salt stress might be mobilized from plastidial MGDG. However, mechanistic evidence or proof for such transport is still missing. Therefore, we focused on a possible coordinated action of plastidial and extraplastidial membranes.

Plant salt tolerance can in fact be influenced by the degree of lipid unsaturation both in the plasma membrane and in thylakoids. For instance, halophytes display high levels of unsaturated fatty acids in MGDG and PG, which are related to the protection of PSI and PSII, leading to reduced PSII inhibition upon exposure to salt and an overall improved salt stress tolerance (Meng et al. 2018; Sui and Han 2014). In the cyanobacterium *Synechocystis sp*. a high content of unsaturated fatty acids in membrane lipids was beneficial for the performance of the photosynthetic machinery during salt stress and had a positive effect on Na^+^/H^+^-antiporters (Allakhverdiev et al. 2001). Similarly, the functionality of plastidial or extraplastidial, ER-associated FADs influences salt tolerance in Arabidopsis (Im et al. 2002; Zhang et al. 2012; Zhang et al. 2009).

There are definite links between plastidial and extraplastidial fatty acid unsaturation. Arabidopsis mutant analysis shows that plastidial or ER-localized FADs can functionally compensate for each other despite their localization in different cellular compartments (Kunst et al. 1988, 1989; McConn and Browse 1996, 1998; McCourt et al. 1987). The underlying transport of unsaturated lipid intermediates can occur both in plastid-ER direction and in ER-plastid direction (Browse and Somerville 1991). It is our working hypothesis that such membrane lipid remodeling based on the inter-organellar exchange of lipid metabolites occurs also during acclimation of membranes to stress conditions, such as salt treatment. In this context, plastidial MGDG might serve as an abundant source of polyunsaturated fatty acids. To date, the underlying transfer mechanisms are still not understood at the molecular level. In particular, it remains unclear how MGDG as a structural thylakoid lipid involved in photosystem assembly and xanthophyll cycling might be mobilized, possibly in a stress-dependent manner, and what the consequences are for thylakoid physiology.

Here we show that mild salt treatment of Arabidopsis seedlings results in increased abundance of both PC and MGDG, and analyze consequences for photosynthetic performance. Similar increases in PC and MGDG are observed in the Arabidopsis *fad3* mutant, for which plastid-ER-directed export of linolenic acid-containing intermediates is well established. A systematic comparison of photosynthetic parameters between salt stressed seedlings and the *fad3* mutant reveals that modulated MGDG abundance and/or fatty acid composition can impair thylakoid function not globally but in the *fad3* mutant has an unexpectedly specific effect on NPQ and the light-driven XC.

## Results

### Mild salt stress of two-week-old Arabidopsis seedlings results in an increased abundance of MGDG and PC

Experiments described in this work were initially aimed at elucidating a possible stress-induced mobilization of MGDG-derived lipid intermediates from the plastid to extraplastidial membranes. To assess changes in plant membrane lipid composition upon salt treatment, a mild salt stress of 50 mM NaCl was applied. This experimental setup was chosen to enable observations within the range of physiological responsiveness of the plants and without causing irreversible damage to the plants. Transferring two-week-old Arabidopsis seedlings to solid ½ MS media containing 50 mM NaCl yielded reproducible effects, and plants were macroscopically not affected by the salt treatment over a recorded 48 h-period of exposure (Fig. 1 A). The salt stress applied was perceived by the plants, as evident from the induction of the marker genes *dehydration-responsive element-binding protein 2A* (*DREB2A*) (Liu et al. 1998; Morimoto et al. 2013) and *responsive to desiccation 29B* (*RD29B*) (Msanne et al. 2011) (Supplemental Fig. 1). Besides known salt stress markers, the transcript abundance of fatty acid desaturase genes *FAD2* and *FAD3* as well as that of *MGDG synthase 1* (*MGD1*) (Kobayashi et al. 2009) were analyzed as controls and found to not change upon salt treatment (Supplemental Fig. 1).

Two-week-old Arabidopsis seedlings treated with 50 mM NaCl were harvested, total lipids were extracted after different times of treatment, and the abundance of PC and MGDG was determined by combined TLC and gas-chromatography (Fig. 1 B). The main fatty acids esterified to PC were 16:0, 18:2^Δ9,12^ and 18:3^Δ9,12,15^, with smaller amounts of associated 18:0 and 18:1^Δ9^. By contrast, the predominant fatty acids associated with MGDG were 16:3^Δ7,10,13^ and 18:3^Δ9,12,15^, with 16:2^Δ7,10^ and 18:2^Δ9,12^ present in traces. MGDG displayed an overall higher degree of unsaturation than PC, and these observations reflect known features of these lipids. The overall sum abundance of fatty acids associated with each lipid enables lipid quantification. The mass abundance of both PC and MGDG was significantly increased after 2 h of salt treatment (Fig. 1 B), mostly due to a significant increase of lipid species containing 16:0, 18:2^Δ9,12^ and 18:3^Δ9,12,15^ for PC, or containing 18:3^Δ9,12,15^ for MGDG, respectively. Over the 2 h salt treatment the abundance of PC almost doubled, while MGDG increased by approx. 20 %. The latter increase occurred over an MGDG basal abundance that was already approx. four times higher than that of PC and by molar volume was similar to that of PC (Fig. 2 B). When extending the duration of the 50 mM NaCl treatment up to 24 h and 48 h, the amounts of PC and MGDG remained increased (Supplemental Fig. 2). A tendency towards an increasing degree of unsaturation in fatty acids associated with PC or MGDG over 48 h of salt treatment (Supplemental Fig. 2) was not statistically significant. The data indicate that mild salt treatment resulted in changes in membrane lipid composition both in plastidial and in extraplastidial membranes.

**Figure 1.**
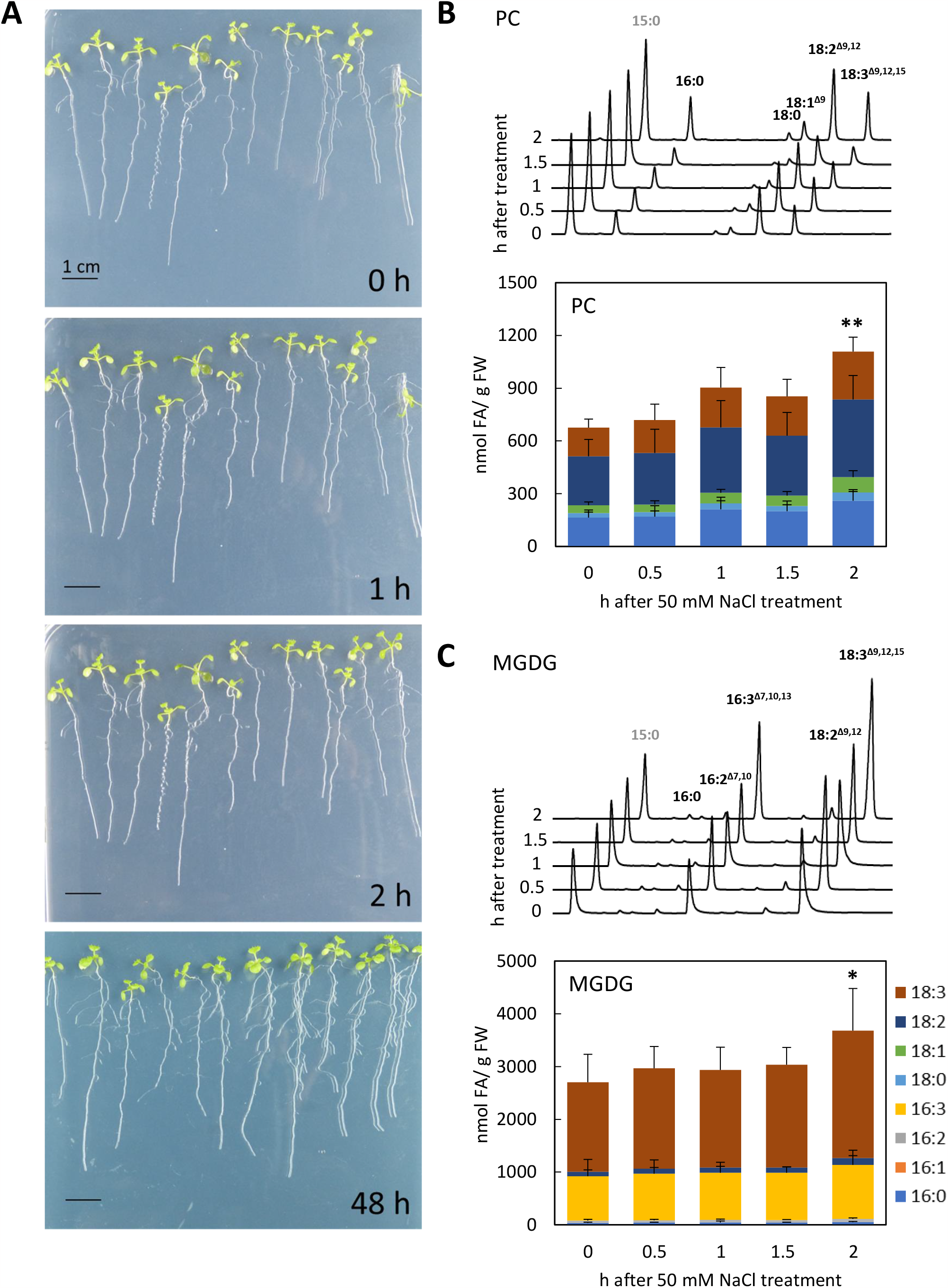
Mild salt treatment results in increased abundance of PC and MGDG. Arabidopsis wild type seedlings were grown for two-weeks on ½ MS solid media and then transferred to ½ MS solid media containing 50 mM NaCl. **A**, The macroscopic appearance of the seedlings was documented after different periods of treatment, as indicated. The images shown are representative for 15 experiments. **B, C**, Green tissue was harvested after the times indicated and the contents of PC and MGDG were analyzed. Lipid extracts were first separated into galacto- and phospholipid fractions by solid phase extraction. Galactolipid and phospholipid classes were then separated by thin-layer-chromatography. Lipid-associated fatty acids were chemically transmethylated and analyzed by gas chromatography. **B**, Upper panel, Representative GC-traces obtained from PC isolated from seedlings exposed to different periods of mild salt treatment. Fatty acid signals as indicated. Lower panel, Molar abundance of PC (whole bar) according to the associated fatty acids (individual bar segments). **C**, Upper panel, Representative GC-traces obtained from MGDG isolated from seedlings exposed to different periods of mild salt treatment. Fatty acid signals as indicated. Lower panel, Molar abundance of MGDG (whole bar) according to the associated fatty acids (individual bar segments). Data represent means ± SD of data from three to four biological experiments, each performed with seven to ten individual samples. Asterisks indicate significant changes of lipid levels compared to time point zero using Student’s T-test (\**P*<0.05; \*\**P*<0.01). Identified fatty acids, blue, 16:0; orange, 16:1^Δ7^; grey, 16:2^Δ7,10^; yellow, 16:3^Δ7,10,13^; light blue, 18:0; green, 18:1^Δ9^; dark blue, 18:2^Δ9,12^; brown, 18:3^Δ9,12,15^.

**Figure 2.**
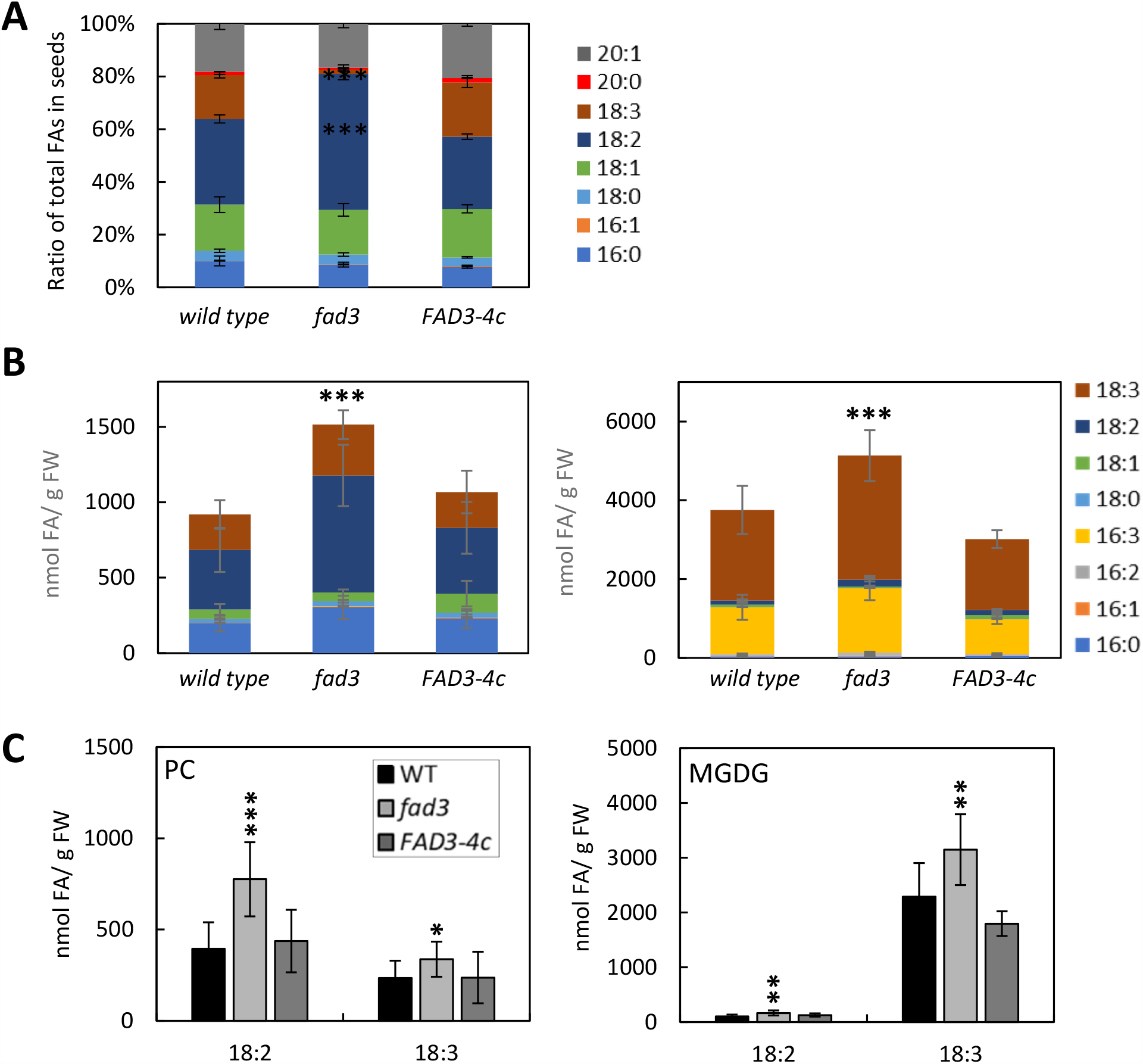
The Arabidopsis fad3 mutant displays increased abundance of PC and MGDG. **A**, Total fatty acids were extracted from seeds of wild type controls, the *fad3* mutant and a complemented *FAD3-4c* line by acidic methanolysis and fatty acids were analyzed by gas chromatography. The proportion of fatty acids in seed oil is given in percent of the total fatty acids. Data represent means ± standard deviations and are from four independent experiments and 6-23 biological replicates. **B, C**, Lipids were extracted from wild type controls, the *fad3* mutant or the complemented *FAD3-4c* line and separated into galacto- and phospholipid fractions by solid phase extraction. Galactolipid and phospholipid classes were then separated by thin-layer-chromatography and PC and MGDG were isolated. Lipid-associated fatty acids were chemically transmethylated and analyzed by gas chromatography. **B**, Molar abundance of PC or MGDG (whole bars), respectively, according to the associated fatty acids (individual bar segments). **C**, Proportion of 18:2 and 18:3 associated with PC and MGDG from wild type controls, *fad3*, or the complemented *FAD3-4c* line, as indicated. Data represent means ± standard deviation from three to five independent experiments, each with 6-12 biological replicates. Asterisks indicate significant changes compared to wild type, using a Student’s T-test (\**P*<0.05; \*\**P*<0.01; \*\*\**P*<0.001).Identified fatty acids, blue, 16:0; orange, 16:1^Δ7^/16:1^Δ9^; grey, 16:2^Δ7,10^; yellow, 16:3^Δ7,10,13^; light blue, 18:0; green, 18:1^Δ9^/ 18:1^Δ11^; dark blue, 18:2^Δ9,12^; brown, 18:3^Δ9,12,15^; red, 20:0; dark grey, 20:1^Δ11^.

### The Arabidopsis *fad3* mutant displays increased abundance of MGDG and PC in green tissues, with wild-type levels of PC-associated linoleic acid

To test whether increased lipid levels in plastids and in extraplastidial membranes were a feature of inter-organellar exchange of lipid intermediates, we analyzed the abundance and fatty acid composition of MGDG and PC in the Arabidopsis *fad3* mutant. This mutant is defective in the formation of linolenic acid at the ER, and in green tissues the defect is metabolically compensated by mobilizing linolenic acid-containing intermediates based on plastidial FAD7/FAD8-mediated linolenic acid formation (Browse and Somerville 1991). Since the effect of the *fad3* mutation was originally described in the context of seed oil accumulation, fatty acids were first extracted from seeds of Arabidopsis wild type controls, the *fad3* mutant and from an Arabidopsis line ectopically expressing a fluorescence-tagged variant of FAD3 under the cauliflower mosaic virus 35S promoter in the *fad3* mutant background (*FAD3-4c*). The predominant fatty acids in seed oil of wild type Arabidopsis were the monounsaturated fatty acids 18:1^Δ9^ and 20:1^Δ11^ (gondoic acid), characteristic for seeds, and the polyunsaturated fatty acids, 18:2^Δ9,12^ and 18:3^Δ9,12,15^ (Fig. 2 A). In seeds of the *fad3* mutant, 18:3^Δ9,12,15^ was reduced compared to wild type controls, concomitant with an increased proportion of 18:2^Δ9,12^, in line with the original description of the *fad3* mutant based on seed analysis (James and Dooner 1990) and highlighting the notion that plastidial fatty acid unsaturation does not contribute substantially to seed oil composition (Fig. 2 A). In the *FAD3-4c* line, the 18:3^Δ9,12,15^ content in seeds was rescued to levels comparable to that of the wild type controls (Fig. 2 A), indicating functional complementation of the *fad3* mutation. Next, we analyzed green tissue of wild type controls, the *fad3* mutant and the complemented *FAD3-4c* line. In contrast to previous reports, in our experiments *fad3* seedlings displayed some developmental retardation compared to wild type, and the defects continued throughout shoot development, resulting in smaller rosettes and shorter inflorescences (Supplemental Fig. 3). The macroscopic defects were rescued in the complemented line, attributing the phenotype to the genetic lesion in the *FAD3* gene (Supplemental Fig. 3). As to our knowledge this phenotype has not previously been described, it is possible that it manifests only under the short-day conditions used in our experiments, compared to continuous light-conditions described before (Browse et al. 1993). The analysis of fatty acids associated with PC isolated from green tissue indicates the presence of substantial amounts of associated 18:3^Δ9,12,15^ (Fig. 2 B). This pattern verifies the previously reported compensatory mobilization of linolenic acid-containing lipid intermediates from the plastids. Based on mass analysis of associated fatty acids, we additionally observed significantly increased abundance of both PC and MGDG in the *fad3* mutant compared to wild type controls or to the complemented *FAD3-4c* line (Fig. 2 B), resulting mostly from an increased proportion of 18:2^Δ9,12^/18:3^Δ9,12,15^ in both lipids (Fig. 2 B, C). The accumulation of 18:2^Δ9,12^ in PC (Fig. 2 B, C) additionally supports an origin of 18:3^Δ9,12,15^ in the plastid, as 18:3^Δ9,12,15^ was present in PC at the same time as accumulation of its precursor 18:2^Δ9,12^ was evident. It is interesting to note that elimination of FAD3 in the ER and ensuing transport of 18:3^Δ9,12,15^ from the plastid to the ER in the *fad3* mutant is accompanied by an increase in the plastidial lipid MGDG as well as in extraplastidial PC (Fig. 2 B). MGDG in the *fad3* mutant was not only enriched in 18:3^Δ9,12,15^ but also contained an elevated proportion of 16:3^Δ7,10,13^, which indicates a higher rate of MGDG biosynthesis via the prokaryotic pathway (Fig. 2 B). This increase occurred despite unchanged transcript abundance of the *MGD1* gene (Supplemental Fig. 1 B). The increased basal levels of PC and MGDG and the increased ratios of lipid-associated 18:3^Δ9,12,15^ in the *fad3* mutant (Fig. 2 B) resembled patterns observed in wild type seedlings after 2 h of salt stress (cf. Fig. 1 B). As the increased lipid levels in the *fad3* mutant are likely a reflection of membrane adjustments during the export of 18:3^Δ9,12,15^ mobilized from MGDG in the plastid to extraplastidial membranes, it is possible that the increased MGDG levels observed upon salt treatment may also indicate such inter-organellar exchange.

**Figure 3.**
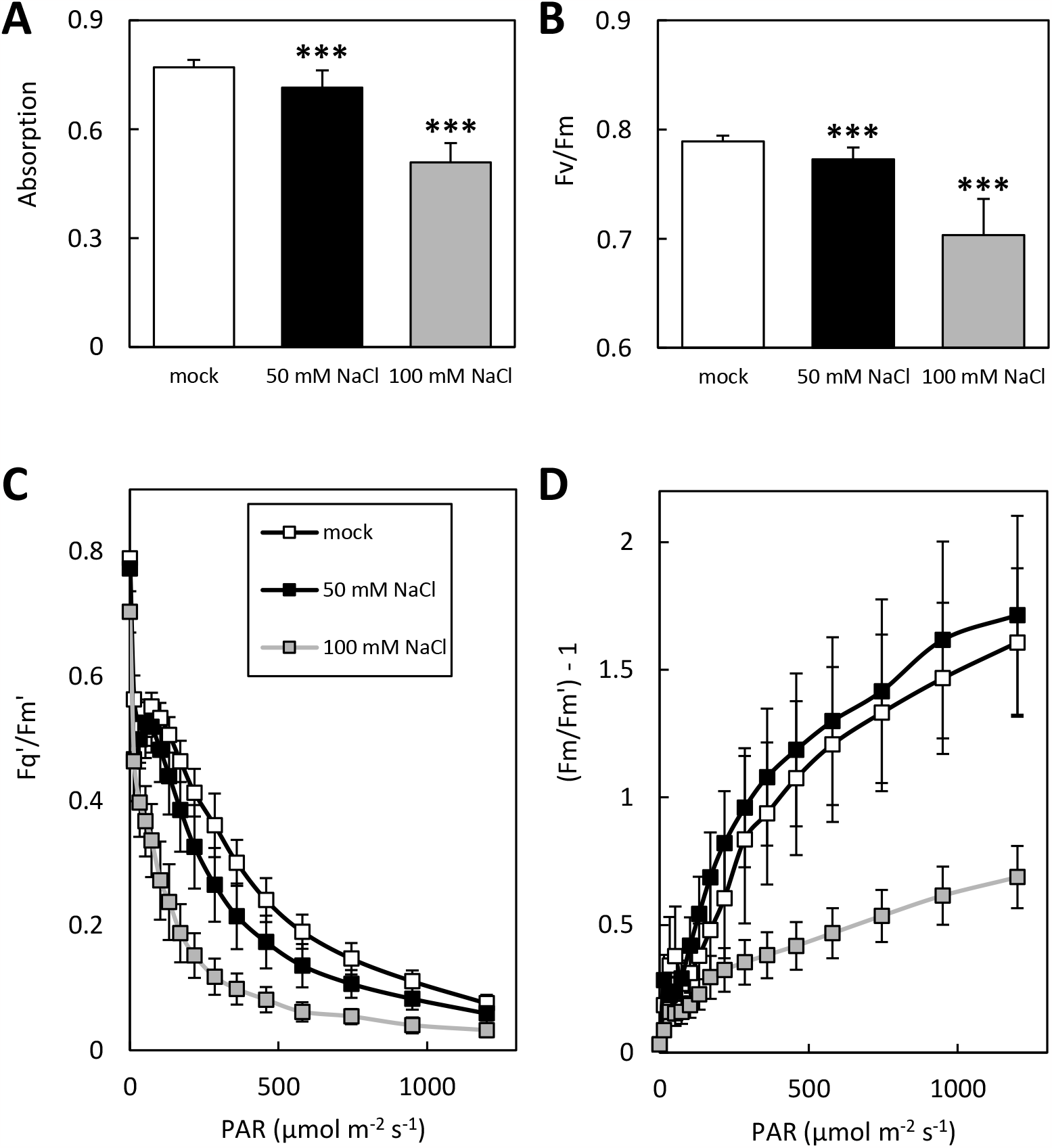
Mild salt treatment results in reduced photosynthetic performance and NPQ. First true leaves of two-week-old wild type seedlings grown on ½ MS solid media without added salt (mock, white), with 50 mM NaCl (black) or with 100 mM NaCl (grey) were harvested and placed on ½ MS agar plates. Leaves were dark-adapted for 5 min before exposure to an 800 ms saturation pulse with a white light intensity of 2400 μmol m^−2^ s^−1^. Subsequently, various photosynthetic parameters were analyzed by IMAGING-PAM. **A**, Absorptivity. **B**, Maximum quantum efficiency of PSII (Fv and Fm). **C, D**, IMAGING PAM was used to determine responses to photosynthetically active radiation (PAR, in µmol m^−2^ s^−1^) increasing in 15 steps of 30 s duration from 14 to 1200 PAR. The recorded parameters Fm, Fm’ and Fq’ were used to calculate the effective quantum efficiency of PSII (**C**) and the non-photochemical quenching (NPQ) at each light intensity (**D**). Data are means ± standard deviations of two independent experiments representing four-five biological replicates. At least five leaves were measured for each biological replicate. Asterisks show significant changes compared to the mock controls according to a Student’s T-test (\**P*<0.05; \*\**P*<0.01; \*\*\**P*<0.001).

To test whether the increased levels of PC or MGDG in the *fad3* mutant were related to lipid increases observed in wild type seedlings upon salt treatments (cf. Fig. 1), relative lipid levels were analyzed upon salt treatment of wild type controls, the *fad3* mutant and the complemented *FAD3-4c* line. While wild type controls displayed an increase in both PC and MGDG after the 2 h period of salt treatment, no such increase was observed in salt treated *fad3* mutants (Supplemental Fig. 4 A), suggesting that the salt-induced mechanism for increasing the lipid levels was not functional in the *fad3* mutant, or was already at capacity with the already-high basal levels of both lipids. The lipid analyses showed similar changes in membrane composition in extraplastidial membranes (PC) or in plastids (MGDG) for wild type seedlings treated for 2 h with 50 mM salt and in the *fad3* mutant. It is our working hypothesis that the elevated lipid levels are related to the inter-organellar mobilization of MGDG-derived lipid intermediates in both cases. To further elucidate possible physiological consequences of the changes in membrane lipid composition, we next focused on the effects of altered MGDG levels, because MGDG is a well-established component of thylakoids, a structural element of photosystem assembly and a key factor of Vx de-epoxidation in the XC. The highly unsaturated MGDG with close to six double bonds per lipid (cf. Supplemental Fig. 2 A) mediates particular membrane phase-order and flexibility in the thylakoids that are biophysically required for the photosynthetic machinery (Hölzl and Dörmann 2019). Therefore, it was tested whether changes in MGDG abundance found in salt-treated wild type seedlings or in *fad3* mutant seedlings had an effect on thylakoid function and photosynthetic capacity using IMAGING-PAM (Figs. 3-6).

**Figure 4.**
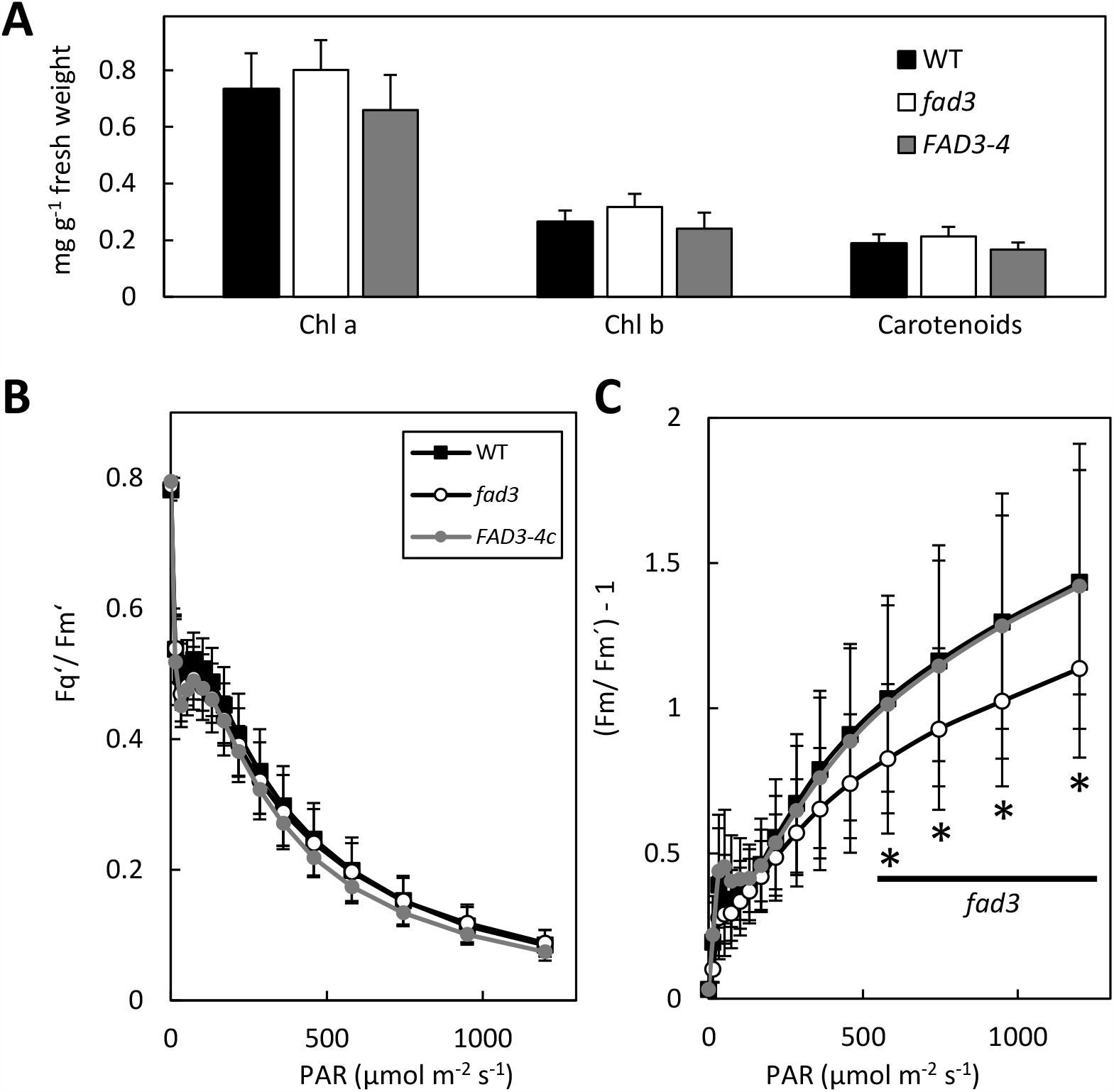
The Arabidopsis *fad3* mutant displays reduced NPQ under high light conditions. Pigments and photosynthetic performance of fad3 mutants were analyzed in comparison to wild type controls and the complemented *FAD3-4c* line. **A**, Pigments were extracted from green tissues of wild type controls, *fad3* mutants or the complemented *FAD3-4c* line using 80 % (v/v) of acetone. Pigments were determined spectrophotometrically by measuring the absorbance at 663, 647 and 470 nm according to (Lichtenthaler and Buschmann 2001), using fresh weight as a reference. Date are means ± standard deviations of two to four independent experiments, representing six to ten biological replicates. Asterisks indicate significant changes compared to WT results using the Student’s T-test (\**P*<0.05; \*\**P*<0.01; \*\*\**P*<0.001). Plant lines: blue, WT; grey, *fad3*; green, *fad78*; dark blue, *fa378*; dark grey, *FAD3-4c*. **B, C**, First true leaves of wild type, fad3 mutants and *FAD3-4c* were harvested and placed on ½ MS agar plates. Leaves were dark-adapted for 5 min before exposure to an 800 ms saturation pulse with a white light intensity of 2400 μmol m^−2^ s^−1^. Subsequently, IMAGING PAM was used to determine responses to photosynthetically active radiation (PAR, in µmol m^−2^ s^−1^) increasing in 15 steps of 30 s duration from 14 to 1200 PAR. The recorded parameters Fm, Fm’ and Fq’ were used to calculate the effective quantum efficiency of PSII (**B**) and the non-photochemical quenching (NPQ) at each light intensity (**C**). Data are means ± standard deviations of two independent experiments representing four-five biological replicates. At least five leaves were measured for each biological replicate. Asterisks show significant changes compared to the mock controls according to a Student’s T-test (\**P*<0.05).

### Salt application has a dose-dependent effect on photosynthesis

Photosynthetic parameters were first determined for two-week-old Arabidopsis wild type seedlings grown on ½ MS solid media (mock) or on the same media with added 50 mM or 100 mM of NaCl (Fig. 3). Absorptivity (Fig. 3 A) is a measure of the fraction of incident red-light which is absorbed by a leaf sample. While the absorptivity values determined by PAM-imaging of leaves do not display the exact concentration of any chlorophyll species, the reading can give an indication of the amount of chlorophyll-absorbed photoactive radiation (PAR) (Fig. 3 A). The mild salt treatment of the plants resulted in gradually decreasing absorptivity with 50 mM or 100 mM NaCl, compared to mock treated controls (Fig. 3 A). Similarly, treatment of seedlings with 50 mM or 100 mM NaCl significantly decreased the maximum quantum efficiency of PSII (Fv/ Fm) (Fig. 3 B). The effective quantum yield of PSII (Fig. 3 C) was also impaired by salt treatment, and plants treated with 50 mM salt displayed reduced effective quantum yields of PSII at normal and higher light intensities compared to mock treated plants, whereas plants treated with 100 mM NaCl exhibited significantly decreased effective PSII quantum yields at all light intensities (Fig. 3 C). An inhibitory effect of the salt treatment was also reflected in a slow and limited increase in NPQ, which was significantly reduced at both low and high light conditions in salt-treated plants compared to mock controls (Fig. 3 D). Energy dissipation at low light intensities was significantly reduced upon treatment with 50 mM NaCl or with 100 mM compared to mock treatment, whereas dissipation was significantly increased at normal intensities and had the same level as in mock-treated plants at high light exposure (Fig. 3 D). Overall, seedlings exposed to salt displayed a dose-dependent impact on photosynthetic performance, in line with previous reports (Johnson and Stepien 2016; Stepien and Johnson 2009, 2018).

### Photosynthetic performance is unaffected, but NPQ is reduced in the Arabidopsis *fad3* mutant

Photosynthetic parameters were next recorded for the *fad3* mutant. First, total pigments were analyzed, and the *fad3* mutant displayed a significantly increased concentration of Chl b compared to wild type controls or the *FAD3-4c* line (Fig. 4 A). When analyzed by IMAGING-PAM, the *fad3* mutant and complemented *FAD3-4c* seedlings both showed effective PSII quantum yields similar to wild type controls at all light intensities (Fig. 4 B). However, when the NPQ was analyzed, the *fad3* mutant displayed significantly lower NPQ values than the wild type when exposed to high light (Fig.4 C), whereas the complemented *FAD3-4c* line showed a comparable energy dissipation to the wild type controls under all light conditions (Fig. 4 C). The analysis of photosynthetic performance indicates that the altered abundance and fatty acid composition of MGDG in the *fad3* mutant was accompanied by reduced photoprotection under high light, with no substantial effect on photosynthetic performance.

### The xanthophyll cycle is inhibited during illumination of the Arabidopsis *fad3* mutant

To more specifically address how a change in thylakoid lipid composition might influence NPQ, we focused on an analysis of the xanthophyll cycle (XC), which contributes substantially to the dissipation of excess light energy as heat. Acidification of the thylakoid lumen by the establishment of the light-driven proton gradient activates the enzyme Vx to Ax and further to Zx, thereby dissipating excess excitation energy (Niyogi et al. 1998). In addition to Vx de-epoxidation, the low luminal pH leads to protonation of the PSII subunit S (PsbS), which belongs to the LHC family of proteins. Protonated PsbS is supposed to play a role in the reorganization of PSII-LHCII super complexes during changes in the light conditions, thereby promoting the conversion of excess excitation energy into harmless heat (Li et al. 2000; Sacharz et al. 2017). Although these processes cannot be truly separated from each other, it has been suggested that the XC is additionally involved in a quenching mechanism that requires a longer induction time and also displays a longer relaxation time after high light illumination (Nilkens et al. 2010). In contrast to the faster quenching processes involving PsbS and Zx, which have been termed high-energy state quanching (qE) the slower Zx-dependent NPQ is referred to as qZ.

Since all NPQ processes occur within thylakoid membranes, it appears reasonable to assume that thylakoid lipids might influence their physiological efficiency. In particular MGDG, which interacts with the LHCII and is additionally required for the operation of the XC within a specialized thylakoid membrane domain, is thought to influence NPQ (Goss and Latowski 2020; Simidjiev et al. 1998; Vieler et al. 2008). The close link between VDE activity and the lipid composition of the thylakoid membrane suggested that changes in the amount and fatty acid composition of MGDG in the *fad3* mutant might affect the efficiency of xanthophyll cycling and thus the level of photoprotection in the mutant in comparison to wild type. To test whether Vx de-epoxidation was perturbed in the *fad3* mutant, wild type or *fad3* seedlings were exposed to different light intensities by the IMAGING-PAM and subsequently used for the extraction and quantitative analysis of leaf pigments by HPLC (Fig. 5). The pigment analysis showed minor differences in the content of neoxanthin (Nx) and lutein (L) between the wild type and the *fad3* mutant (Fig. 5 A). The HPLC data furthermore confirmed the spectral analyses, again showing a higher Chl b concentration in the *fad3* mutant than in the wild type controls (cf. Fig. 4 A). The data on XC-pigment conversion at the different light intensities from 0 to 1200 PAR are shown as the de-epoxidation state (DES) of the XC in wild type and the *fad3* mutant. The DES, which depicts the percentage of de-epoxidized XC pigments, i.e. Ax and Zx, of the complete XC pigment pool, serves as a measure for the efficiency of the conversion of Vx to Ax or Zx at a given light intensity (Fig. 5 B). While the DES increased in wild type plants under normal light conditions (170 PAR), in *fad3* plants a decrease of the DES at this light intensity compared to dark-adapted leaves was observed (Fig. 5 B). In the wild type leaves the DES increased further until high light conditions of 950 PAR were reached. The *fad3* mutant was also characterized by a light intensity-dependent increase of the DES. However, the DES values were always substantially lower than those in the wild type controls (Fig. 5 B). For a better illustration, the proportion of the individual XC pigments of the complete XC pigment pool are depicted in Fig. 5 C for the wild type control and for the *fad3* mutant. While the wild type control displayed a visible increase of Ax and Zx at the expense of Vx with exposure to increasing light intensities, the contents of the XC pigments changed only marginally in the *fad3* mutant, apart from a slight increase in Zx at 950 PAR (Fig. 5 C). These observations suggest a possible XC defect in the *fad3* mutant at the level of VDE function.

**Figure 5.**
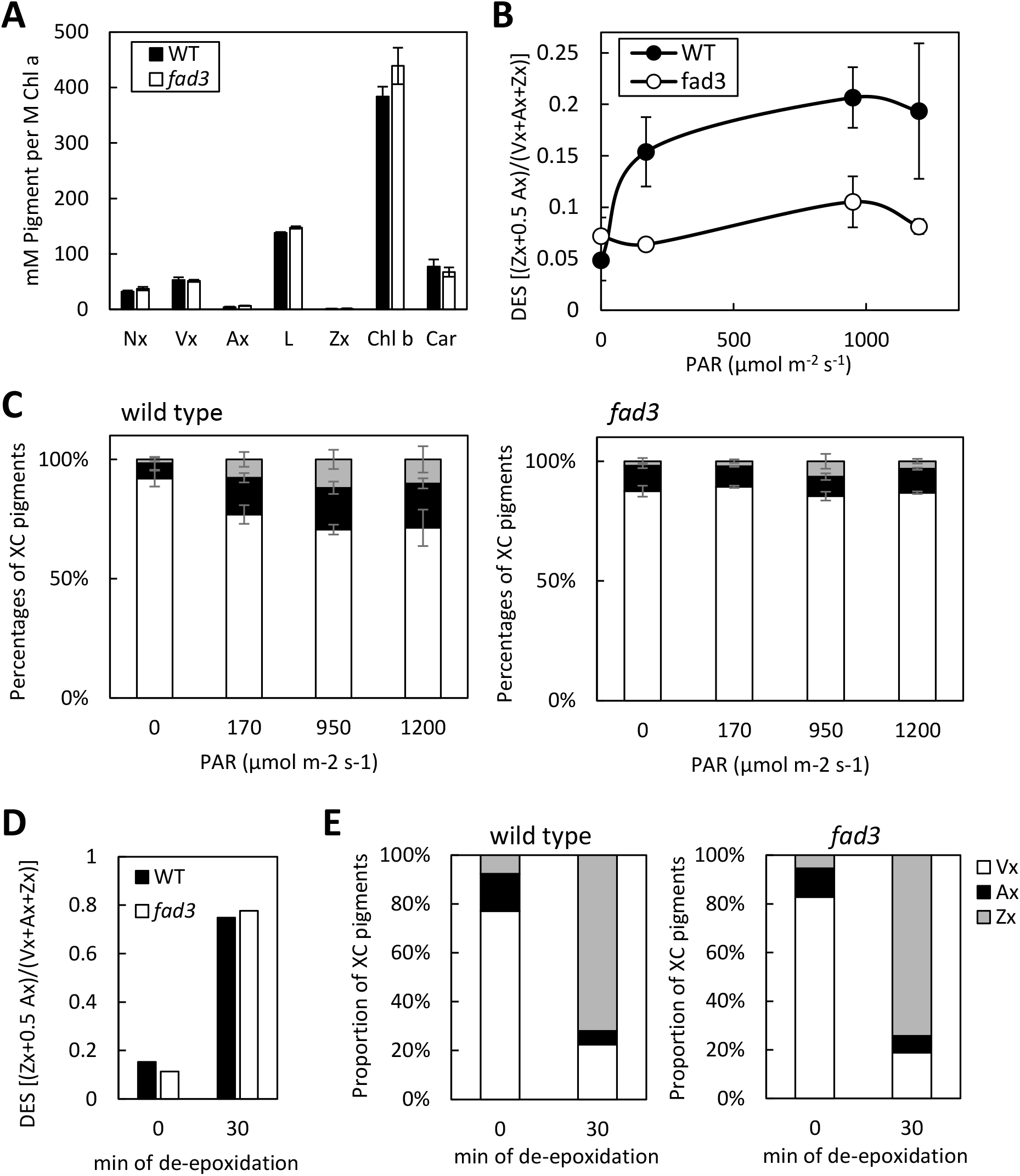
The Arabidopsis *fad3* mutant displays a reduced de-epoxidation of xanthophyll cycle-pigments under high light conditions. First true leaves of wild type controls or *fad3* mutants were placed on ½ MS solid media and dark-adapted for 5 min before starting with a saturation pulse of 800 ms with a light intensity of 2400 μmol m^−2^ s^−1^. A light curve with photosynthetic active radiation (PAR, in µmol m^−2^ s^−1^) was recorded by IMAGING-PAM using 30 s exposure to 0, 170, 950 and 1200 PAR. After exposure to the different light intensities during the illumination program, xanthophyll cycle-pigments were extracted and analyzed by HPLC. **A**, amounts of neoxanthin (Nx), violaxanthin (Vx), antheraxanthin (Ax), lutein (L), zeaxanthin (Zx), chlorophyll b (Chl b) and beta-carotene (Car) in mM pigments per M of chlorophyll a (Chl a), measured at 0 PAR. **B**, The pigment determinations after exposure to the different PAR intensities were used to calculate the light-dependent de-epoxidation states of the xanthophyll cycle pigment pool (DES). **C**, The proportions of individual XC pigments of the complete xanthophyll cycle pigment pool for different PAR intensities were determined for wild type controls (left panel) or the *fad3* mutant (right panel). Vx (white), Ax (black) and Zx (grey). Data represent means ± standard deviations representing two to three biological replicates. **D, E**, Analysis of Vx de-epoxidation under optimal conditions with intrinsic VDE in isolated thylakoids from leaves of wild type controls and *fad3* mutants. **D**, DES calculated from the concentrations of XC-pigments in thylakoids isolated from wild type controls (black bars) or from *fad3* mutants (white bars), as determined by HPLC before and 30 min after the start of the de-epoxidation reaction with ascorbic acid. **E**, Proportion of the XC pigments violaxanthin (Vx, white), antheraxanthin (Ax, black) and zeaxanthin (Zx, grey) of the total xanthophyll cycle pigment pool before and 30 min after the start of the de-epoxidation reaction in thylakoids from wild type controls or from *fad3* mutants, as indicated. Data are representative for two independent experiments.

The lower DES observed in the *fad3* mutant might be the consequence of a reduced activity of the VDE, a reduced access to its substrate Vx or its co-substrate ascorbate, and/or a less efficient establishment of the lipid structures within the thylakoid membrane that are needed for the de-epoxidation process. To experimentally differentiate between these possibilities, thylakoids were isolated from wild type controls or from *fad3* mutants and used for *in vitro* de-epoxidation assays under optimal reaction conditions for the intrinsic VDE at pH 5.0 and a temperature of 30 °C. Samples were taken prior to the start of the de-epoxidation reaction by the addition of the co-substrate ascorbate (time point 0 min) and after 30 min of the conversion of Vx to Zx. XC pigments were extracted and their concentrations determined by HPLC (Fig. 5 D), the state of the XC pigment pool was again depicted as DES. Under the conditions of the de-epoxidation assays, the wild type control and the *fad3* mutant displayed a strong conversion of Vx to Zx and up to four-fold increases in the DES could be determined after 30 min of the assay, compared to the values at 0 min (Fig. 5 D). Importantly, the DES did not differ in thylakoids from the wild type control or the *fad3* mutant. In both cases Vx was very efficiently converted to Ax and Zx (Fig. 5 D), demonstrating an equally high VDE activity in thylakoid membranes isolated from both wild type and *fad3* seedlings. Thus, the results from the de-epoxidation assays in the dark under optimal conditions indicate that the reduced Vx de-epoxidation in the illuminated *fad3* mutant leaves was not caused by differences in the VDE activity itself. The different thylakoid lipid and fatty acid composition might have had an impact on substrate/co-substrate availability during illumination or even on light-driven establishment of the special lipid structures that are needed for the efficient conversion of Vx to Ax and Zx.

## Discussion

The lipid composition of membranes is an important element of plant responses to stress conditions, both to maintain the integrity of membranes and to enable the function of integral membrane protein complexes. We report on simultaneous increases of MGDG and PC abundance within 2 h of mild salt treatment (Fig. 1) and hypothesize that these increases are mechanistically related to a postulated plastid-ER-directed exchange of lipid intermediates. The hypothesis is addressed by comparing the effects of salt treatment on membrane lipids and thylakoid physiology with those in the Arabidopsis *fad3* mutant, for which a plastid-ER-directed exchange of linolenic acid-containing intermediates is known (Browse and Somerville 1991). Interestingly, the Arabidopsis *fad3* mutant also displayed an increased abundance of MGDG and PC without any salt treatment, thus displaying a lipid pattern similar to that observed for salt-treated wild type Arabidopsis seedlings (Fig. 2). As no further changes in lipid abundance were observed after exposure of *fad3* mutants to salt (Supplemental Fig. 2), it is possible that the increased lipid levels are related to inter-organellar lipid mobilization in both metabolic scenarios.

Besides its well known role as a structural element of thylakoid membranes with an important function in photosynthetic reactions, MGDG serves as a reservoir for polyunsaturated fatty acids to be mobilized from plastids under stress conditions. The contribution of linoleic acid and hexadecatrienoic acid for the wound-induction of jasmonic acid biosynthesis is a well-studied example for stress-induced mobilization of MGDG-derived lipid intermediates. However, jasmonic acid production occurs at a small molar scale compared to the over-abundance of MGDG (Mosblech et al. 2009; Wasternack and Feussner 2018), so this process will not result in a detectable change in the abundance of MGDG. By contrast, previous observations using different stresses, such as salt treatment (König et al. 2008; König et al. 2007) or wounding (Mosblech et al. 2008), indicate more massive changes in MGDG that suggest stress-induced mobilization of MGDG for an alternative metabolic purpose. The increases in MGDG and PC observed even upon mild salt treatment were still substantial (Fig. 1), roughly representing a doubling in the abundance in PC and an approx. 20% increase in MGDG, the most highly abundant membrane lipid in plants. The inter-organellar mobilization of 18:3^Δ9,12,15^-containing lipid intermediates in the *fad3* mutant has been known for decades (Browse et al. 1993; Browse and Somerville 1991), and the finding that especially MGDG levels were correspondingly altered suggests an inter-organellar influence on the composition and organization of plastidial membranes, including thylakoids.

The biosynthetis of MGDG and PC increased upon stress treatment in addition to the already-high abundance of these lipids, suggesting that the newly-formed lipids might have a metabolic fate different from that of the MGDG and PC present prior to stimulation, which might not have been available for mobilization. As plants have evolved in the presence of varying environmental conditions, it appears reasonable to assume that the metabolic availability of mobilizable MGDG might pose an evolutionary advantage. In line with this thought, overexpression of the MGDG-synthase OsMGD enhanced salt tolerance in rice, due to a higher protection of the photosynthetic apparatus and a stabilized thylakoid structure (Guo et al. 2019; Sui and Han 2014; Wang et al. 2014). In previous long-term salt experiments, the abundance of membrane lipids was found to differ between Arabidopsis and its salt-tolerant relative, *Thellungiella halophila*, and it was proposed that membrane lipid composition is part of the mechanisms conferring salt tolerance to halophytes (Sui and Han 2014). In *T. halophila*, the amount of 18:3^Δ9,12,15^-containing MGDG species increased at the expense of 16:0-containing MGDG species when treated with 100, 200 and 300 mM NaCl. While in Arabidopsis the degree of unsaturation of MGDG did not change when the plants were exposed to 100 mM NaCl, exposure to 200 mM NaCl resulted in a significant decrease in the degree of unsaturation, which was proposed to be the result of MGDG degradation due to induced senescence mechanisms (Sui and Han 2014). It appears possible that thylakoids have an intrinsic capacity to form MGDG-based metabolites for alternative use, and that the abundance of MGDG might be related to plant stress tolerance. It is interesting to note that changes in MGDG and PC observed upon mild salt treatment differed from the dramatic transient decreases in these lipids reported in earlier studies (König et al. 2007; Mosblech et al. 2008). While the experimental conditions used here differed substantially from those used in earlier reports, the differences in observed lipid patterns might be explained in part by the different intensity of the stress applied (e.g., 50 mM NaCl vs. 400 mM NaCl in (König et al. 2007)). Importantly, even the earlier studies reported increased formation of MGDG and PC, because transient reductions in these lipids were followed by increases to base levels, respectively (König et al. 2007; Mosblech et al. 2008). Thus, it can be postulated that (re-) synthesis of MGDG might be part of the stress responses observed in these cases.

It is currently unclear whether alternative roles of MGDG as a functional element of thylakoids or as a mobilizable resource might occur at mutual expense. So far, only little is known about the lipid nano-organization in thylakoids, and we are just beginning to understand the role of MGDG and other lipid species in thylakoid function. Our data show that the altered plastidial MGDG abundance in the *fad3* mutant on thylakoid function did not exert a global effect on photosynthesis, but instead had a limited effect only on NPQ and specifically the XC under high light intensities (Fig. 5), which involves proteins peripherally associated with the photosystem core complex, i.e. LHCII, or enzymes which only transiently bind to the thylakoid membrane, such as VDE (Goss and Latowski 2020; Li et al. 2000). This observation suggests that the evident mobilization of plastidial MGDG in the *fad3* mutant affects only part of the MGDG present in the thylakoids. As MGDG molecules tightly bound as structural elements of the protein-pigment complexes might be difficult to dislodge and mobilize for the export of MGDG-derived intermediates, MGDG molecules that are less tightly associated with PSI or PSII, or MGDG molecules that do not directly interact with proteins might be preferentially mobilized. The previous observation that MGDG is substantially degraded in Arabidopsis leaves upon severe (400 mM) salt treatment (König et al. 2007) suggests that the postulated export of MGDG-derived lipid intermediates from the plastid might be prioritized over maintaining thylakoid function. An intense stress dosage used for instance by König and coworkers might rapidly deplete MGDG available for mobilization, resulting in additional (and undesired) effects also on MGDG that is physiologically active within the membrane, with possibly catastrophic consequences for thylakoid function. By contrast, the mild stress applied here (Fig. 1) might promote enhanced biosynthesis of MGDG and a limited mobilization of MGDG-derived metabolites towards extraplastidial membranes still within the physiological capacity of the plants and without impairing growth or posing damage to thylakoids.

The altered membrane lipid composition observed either upon salt stress or in the *fad3* mutant nonetheless had an impact on membrane function. Effects of the mild salt stress on plastid physiology of wild type seedlings reflected previously reported salt-mediated inhibition of photosynthesis (Johnson and Stepien 2016; Stepien and Johnson 2009, 2018). While the intensity of the salt treatment applied was sufficiently low as not to impair plant growth (Fig. 1 A), the expression analysis of marker transcripts (Supplemental Fig. 1) indicated that the stress applied was sufficient to induce salt responses, and photosynthetic parameters were affected (Fig. 3). The consequences of salt stress were a decrease in the effective quantum yield of PSII and an altered NPQ (Fig. 3). By comparison, seedlings grown on 100 mM NaCl generally showed a greater reduction in the photosynthetic efficiency of PSII than those exposed to 50 mM NaCl, probably due to inactivation of PSII, as also indicated by a low optimal quantum efficiency of PSII (Fig. 3) (Maxwell and Johnson 2000). The effects of 100 mM NaCl resulted in a lower NPQ, even though the excitation energy was not used efficiently for photochemistry (Fig. 3). A possible explanation for this apparent “loss of electrons” is that treatment with 100 mM salt might have caused partial disruption of the thylakoid membrane, and thylakoid leakage might have interfered with the build-up of the proton gradient that leads to PsbS protonation and controls the activation of VDE, and thus the formation of Zx as an NPQ component. As VDE activity in thylakoids isolated from wild type controls or *fad3* mutants did not differ during darkness under optimal conditions (Fig. 5 D, E), the effects of an altered MGDG formation on Vx de-epoxidation in the *fad3* mutant have to be light-dependent. In this regard, the comparable effective quantum yields between wild type and the *fad3* mutant (Fig. 4) argue against the role of an impaired proton gradient for the decreased NPQ and DES values that were observed during illumination in the mutant. It is possible that the altered thylakoid lipid composition in the *fad3* mutant impaired the access of VDE to its substrate Vx or its co-substrate ascorbate during illumination. It can also be hypothesized that the light-dependent establishment of inverted hexagonal MGDG phases, which act as attraction sites for the transient binding of VDE to the thylakoid membrane (Garab et al. 2016), is somewhat hindered in the *fad3* mutant with its altered lipid and fatty acid composition. In addition to Zx formation, the processes contributing to NPQ also involve the special PSII antenna subunit, the PsbS protein. PsbS can associate with the PSII-LHCII super complexes, resulting in a rearrangement of the antenna complexes within the thylakoids to form heat-dissipating centers (Kaiser et al. 2019). Besides the effect on Vx de-epoxidation, the mobilization of MGDG in the *fad3* mutant (and possibly also upon salt treatment) might impair the lateral diffusion of proteins at the PSII periphery, such as PsbS, that are required to convert PSII from a light-harvesting into a heat-dissipating state and *vice versa*. Such an impairment of the diffusion of special proteins might account for the observed specific effect of the altered lipid composition of the *fad3* mutant on NPQ.

Taken together, the data presented show that defective ER-based membrane lipid unsaturation in the *fad3* mutant is compensated by the export of polyunsaturated fatty acids from the plastid (Fig. 2 B), as was previously demonstrated (Browse and Somerville 1991). Importantly, this export is accompanied by a substantial increase in the main thylakoid lipid, MGDG, which occurs in addition to an already high abundance of this lipid. An equivalent increase in MGDG was also observed upon mild salt treatment (Fig. 1 B), for which export of polyunsaturated fatty acids from the plastids to extraplastidial membranes can be proposed but has not been demonstrated (König et al. 2007). The data suggest that an extraplastidial requirement for lipid intermediates can promote the formation of MGDG to enable the export of polyunsaturated fatty acids from the plastid. The mobilization of MGDG will affect the photosynthetic capacity of thylakoids, as evident in the *fad3* mutants and also in seedlings exposed to increasing intensities of salt stress. The rather specific effects of compensatory MGDG mobilization in the *fad3* mutant on the XC might be a consequence of a better accessibility of membrane domains where the de-epoxidation of Vx to Zx is taking place, so MGDG in these domains might be prone to be recruited for export, rather than MGDG in areas occupied more tightly by the protein-dense photosystems. The effects observed might, thus, provide a limited and circumstantial insight into thylakoid membrane organization.

## Material and Methods

### cDNA constructs

The cDNA encoding EYFP was amplified using the primer combination 5’-ATGCGGCGCGCCATGGTGAGCAAGGGCGAGGA

-3’/5’-ATGCCTCGAGCTTGTACAGCTCGTCCA-3’ and moved as an AscI/XhoI fragment into the vector pEntryA, creating *pEntryA-EYFP*. The coding sequence for FAD3 was amplified using the primer combination 5’-

ATGCCTCGAGATGGTTGTTGCTATGGACCA-3’/5’-ATGCGCGGCCGCTTAATTGATTTTAGATTTGT-3’ and moved as an XhoI/NotI fragment in frame with the EYFP coding sequence into *pEntryA-EYFP*, creating *pEntryA-EYFP-FAD3*. For plant transformation driven by the 35S-promoter, the EYFP-FAD3 coding sequence was moved from *pEntryA-EYFP-FAD3* into the plasmid *pCAMBIA3300*.*0GS* using Gateway technology according to manufacturer’s instructions.

### Plant material and growth conditions

Arabidopsis (*Arabidopsis thaliana*) seeds of Columbia-0 (Col-0) wild-type and mutant lines *fad3-2* (*fad3*) or *FAD3-4c* were surface sterilized using 6 % (w/v) NaOCl/ 0.001% Triton X-100 and washed six times with sterile water before incubation in 0.1 % (w/v) agarose at 4 °C for two days. Seeds were sown on plates containing 1/2 Murashige and Skoog media (1/2 MS), 1 % (w/v) sucrose and 0.8 % (w/v) plant agar before being placed vertically for two weeks under short day conditions (8 h illumination of 170–200 µmol m^−2^ sec^−1^, 21 °C; 16 h dark, 18°C). To account for the developmental retardation of the *fad3* mutant, experiments with *fad3* mutants were performed after three weeks of growth when plants were at a developmental stage equivalent to two-week-old wild type plants.

### Arabidopsis transformation

Arabidopsis *fad3* mutants were transformed by floral dipping as previously described (Clough and Bent 1998). The resulting transgenic plants were selected according to herbicide resistance. The *FAD3-4c* line used is in the T3 generation and fully complements the *fad3* mutant defect. The line is representative for 12 independent complemented lines generated.

### Arabidopsis stress treatments

Vertically grown plants were transferred to ½ MS solid media containing 0.05 M or 0.1 M NaCl. After different incubation times plants were photographically documented and harvested for further analysis into 15 ml reaction tubes.The samples contained pools of 13-15 individual plants and were frozen in liquid nitrogen. For measurements of photosynthetic parameters, seedlings were grown horizontally for two weeks on ½ MS solid media under control conditions or with either 0.05 M or 0.1 M NaCl added under short-day conditions (8 h light / 16 h dark, 21 °C/ 18 °C; 170 µmol m^-2^ s^-1^). The first true leaves were harvested off and placed on ½ MS solid media for dark adaptation and subsequent IMAGING-PAM measurements.

### Lipid extraction, fatty acid derivatization and analysis

Plant material was harvested and ground in liquid nitrogen. Lipids were extracted according to (Bligh and Dyer 1959), using 0.15 M NaCl instead of water. From total lipid extracts, galactolipids and phospholipids were enriched by solid phase extraction as previously described (Launhardt et al. 2021). Lipids from each fraction were separated by thin-layer-chromatography using developing solvents containing acetone:toluene:water (90:30:7, v/v/v) for galactolipids or chloroform:methanol:acetic acid (65:25:8, v/v/v) for phospholipids, respectively. TLC plates were sprayed with 0.02 % (w/v) primuline (acetone:water, 4:1, v/v) and lipids were visualized under UV-light. MGDG and PC were isolated from the plates, the associated fatty acids were chemically transmethylated and analyzed by gas chromatography using a GC-2010 plus GC/FID-system (Shimadzu) equipped with a 30 m x 250 µm DB-23 capillary column (Agilent) as previously described (Launhardt et al. 2021). Extraction and derivatization of Arabidopsis seed total fatty acids was performed with 10 µl of 0.2 M trimethyl sulfonium hydroxide (Butte et al. 1982). The resulting FAMEs were dissolved in 10 μl acetonitrile and analyzed by gas chromatography as described.

### Spectrophotometric pigment analysis

Shoot material (approximately 12 mg) of vertically grown plants was harvested at the end of the dark phase into 2 ml reaction tubes containing six 3 mm glass beads and 200 µl distilled water. Samples were kept in the dark and on ice. After determination of the fresh weight, the material was homogenized at 4 °C for 40 s in a shaking mill at a frequency of 30 motions s^-1^. Afterwards, 800 µl cold acetone was added and the mixture incubated rotating at 4 °C for 40 min in an overhead shaker. Samples were centrifuged for 5 min at 18,500 x g and 4 °C and 800 µl of the supernatant transferred into a single-use cuvette. The absorbance was measured at 663, 647 and 470 nm against 80 % acetone as a blank in an Ultrospec 2100 pro photometer for chlorophyll a, chlorophyll b and carotenoid determination, respectively. Pigment amounts were calculated according to Lichtenthaler and Buschmann (Lichtenthaler and Buschmann 2001) and referenced to the respective fresh weights.

### Thylakoid isolation

Thylakoids were isolated as previously described (Casazza et al. 2001). Green tissue was harvested from approximately 500 horizontally grown seedlings at the end of the dark phase. After incubation in iced water for 30 min, plants were homogenized in 5 ml of a buffer containing 0.4 M sorbitol, 5 mM EDTA, 5 mM EGTA, MgCl_2_, 10 mM NaHCO_3_, 20 mM Tricine (NaOH; pH 8.4) and 0.5 % (w/v) fatty acid free bovine serum albumin by using mortar and pestle and filtered through two layers of cotton gauze. For maximum yield, the gauze was additionally washed with 10 ml of the same buffer. Thylakoids and remaining intact chloroplasts were sedimented at 2,600 x g and 4 °C for 3 min. Sediments were washed by resuspension in 3 ml buffer containing 0.3 M sorbitol, 2.5 mM EDTA, 5 mM MgCl_2_, 10 mM NaHCO_3_, 20 mM HEPES (NaOH; pH 7.6) and 0.5 % (w/v) fatty acid free BSA. The washing step was repeated one more time. To disrupt remaining intact chloroplasts, the sediment was first resuspended in 1 ml of hypotonic buffer containing 2.5 mM EDTA, 5 mM MgCl_2_, 10 mM NaHCO_3_, 20 mM HEPES (pH 7.6) and 0.5 % (w/v) fatty acid free BSA, and the volume was then brought to 5 ml. After centrifugation, the sediment was resuspended in 1 ml of resuspension buffer and the thylakoid preparation was stored on ice in the dark. The amount of isolated thylakoids used for the present experiments was based on the calculated total chlorophyll content and is given as µg total chlorophyll mg^-1^. The total chlorophyll content of the thylakoid preparations was determined by dissolving 30 µl of thylakoid suspension in 5 ml of 80 % acetone. After centrifugation at 4 °C at 2,600 x g for 5 min, 1 ml of the supernatant was used for photometric determination of chlorophyll a and chlorophyll b absorbance at 663 nm and 647 nm, respectively.

### Chlorophyll fluorescence and photosynthetic performance

First true leaves of horizontally grown seedlings were harvested and placed on 1/2 MS media plates. After dark adaption for at least five minutes, fluorescence parameters were detected using an IMAGING-PAM chlorophyll fluorometer that consists of an IMAG-C control unit combined with an IMAG-L LED-Ring-Array and an IMAG-K CCD camera (Heinz Walz, Effeltrich, Germany). The pulse-amplitude-modulated measuring light (470 nm) was applied with an intensity of approx. 0.1 μE m^−2^ s^−1^ and a modulation frequency of 1 Hz. A saturation pulse was used with a duration of 800 ms and an intensity of 2400 μE m^−2^ s^−1^. Minimal fluorescence (Fo) and maximal fluorescence (Fm) parameters were determined at dark adapted leaves to obtain the maximum PSII quantum efficiency (Fv/Fm). A light-curve with 15 steps of photosynthetic active radiation (PAR [in µE m^−2^ s^−1^]), increasing from 14 to 1200 μE m^−2^ s^−1^ with duration intervals of 30 s, was subsequently recorded. The parameters F’ (steady state fluorescence directly before the saturation pulse) and Fm’ (maximal fluorescence during actinic illumination) were determined at the end of each light step and the effective PSII quantum yield (Yield = Fq’/Fm’), non-photochemical quenching (NPQ = (Fm/ Fm’) – 1) and electron transport rate (ETR = 0.5 x Yield x PAR x Abs. [in µequivalents m^−2^ s^−1^]) were calculated. Each individual leaf absorptivity is estimated by Abs. = 1 – R/NIR (R = value of red light (650 nm) remission; NIR = value of near infra-red light (780 nm) remission) according to the manufacturer’s instructions.

### Analysis of xanthophyll cycle pigments

The analysis of the de-epoxidation state of xanthophyll cycle (XC) pigments was performed either during high light illumination of intact leaves, thus assessing the light-driven de-epoxidation of Vx to Zx; or in isolated thylakoids in the dark under optimal conditions for the enzyme VDE, i.e. in a buffer adjusted to the pH-optimum of the enzyme. To assess the light-dependent, intrinsic degrees of Vx de-epoxidation, five leaves of horizontally grown seedlings were collected and placed on ½ MS agar plates. Leaves were exposed to photosynthetic active radiation from an IMAGING-PAM chlorophyll fluorometer. Stepwise increases of radiation from 14 to 1200 µE m^-2^ s^-1^ were used to expose leaves to different doses of radiation. Samples were taken when after the stepwise increase a certain light intensity had been reached (0, 170, 950, 1200 µE m^-2^ s^-1^ of radiation). Leaves were collected in tubes containing glass beads (size 0.85-1.23 mm) and frozen in liquid nitrogen. For the extraction of pigments, leaves were first ground for 1 min at -80 °C in a shaking mill at a frequency of 30 motions s^-1^ and then for a second time after adding 750 µl of chloroform:methanol:ammonia (1:2:0.004, v/v/v). A volume of 750 µl of distilled water was added and the samples were centrifuged at 21,100 x g and 4 °C for 1 min. The lower phase (200 µl) was transferred to a new tube, samples were evaporated in a stream of nitrogen and stored at - 20 °C until they were analyzed by HPLC. To determine Vx de-epoxidation in isolated thylakoids, under optimal conditions, the conversion of Vx to Zx was triggered by incubation in a reaction buffer adjusted to pH 5, and sodium ascorbate, the co-substrate of the de-epoxidation reaction, was added at a concentration of 30 mM.Isolated thylakoids corresponding to final 20 µg total chlorophyll content ml^-1^ were pre-incubated for 5 min in 1 ml of buffer containing 10 mM NaCl, 5 mM MgCl_2_ and 40 mM MES pH 5.0 at 30 °C in the dark. De-epoxidation was initiated by adding a further 4 ml of the same buffer which additionally contained 30 mM ascorbate,followed by the immediate sampling of 750 µl of the mixture at time point 0 min. Reactions were incubated for 30 min at 30 °C in the dark and additional samples were taken at the end of the 30 min incubation period. To stop the de-epoxidation reaction and extract the pigments, 750 µl of sample was injected into 750 µl of chloroform:methanol:ammonia (1:2:0.004, v/v/v), the mixture centrifuged for 1 min at 4 °C at 21,100 x g, and 200 µl of the lower phases were transferred to new tubes. Solvents were evaporated under a stream of nitrogen and the dried pigments kept at -20 °C until they were analyzed by HPLC. For pigment analysis by HPLC, the dried samples were resuspended in 200 µl of a solution containing 0.2 M ammonium acetate (9:1, v/v) and 10 % (v/v) ethyl acetate in methanol and transferred to HPLC injection glass vials. Pigments were separated and analyzed using an UltiMate 3000 HPLC system (Dionex). For each measurement, 100 µl of sample were injected using a WPS-3000SL autosampler (Dionex) into an ET 250/4 Nucleosil 120-5 C18 chromatography column.Pigments were separated in 35 min runs using an elution gradient at a flow rate of 1.2 ml min^-1^ as previously described (Frommolt et al. 2001). For an accurate mix of solvents, an LPG-3400A system (Dionex) was used, consisting of a quaternary low-pressure pump with integrated vacuum degasser. The column thermostat TCC-3000 (Dionex) ensured a constant temperature of 25 °C for the duration of the chromatographic separation. During the whole run the photodiode array detector PDA-(Dionex) 3000 detected eluting pigments in a wavelength range between 350-750 nm and the chromatograms were recorded using Chromeleon software (Dionex). For quantification of pigments, values at 440 nm were used as previously described (Böhme et al.2002). Further absorption spectra were saved for post-processing. Conversion of Vx to Ax and Zx was calculated as the de-epoxidation state (DES) of the xanthophyll pigment pool using the formula DES =(0.5Ax+Zx)/(Vx+Ax+Zx).

### Relative quantification of transcript levels

Total RNA was isolated by the Trizol method and transcribed into cDNA using the RevertAid H Minus First Strand cDNA Synthesis Kit (Thermo Fisher Scientific). Real-time qPCR was performed using the Luna Universal Qpcr Master Mix (New England Biolabs), according to manufacturer’s instructions. Data were normalized to *UBC10* as an internal reference.

## Accession numbers

Gene loci investigated in this study are identified as follows: *DREB2B*, At3g11020; *FAD2*, At3g12120; *FAD3*, At2g29980; *MGD1*, At4g31780; *RD29B*, At5g52310; *UBC10*, At5g53300

## Data Availability

All materials generated in this study are available upon request.

## Funding

This work was supported by the German Research Foundation (DFG, grant 400681449/GRK2498 TP01 to I.H. and 400681449/GRK2498 TP09 to K.H.).

## Acknowledgements

The authors acknowledge helpful discussion by Leander Ehmke (Martin Luther University Halle-Wittenberg).

## Author contributions

Conceptualization, I.H.; methodology, I.H., K.H. and R.G.; investigation, M.M., L.L., O.B., R.G.; data curation, I.H., M.M., R.G.; writing—original draft preparation, I.H., M.M.; writing, review and editing, I.H., M.M., L.L., O.B.,K.H., R.G.; project administration, I.H.; funding acquisition, I.H., K.H. All authors have read and agreed to the submitted version of the manuscript.

## Disclosures

The authors declare no conflict of interest. The funders had no role in the design of the study; in the collection, analyses, or interpretation of data; in the writing of the manuscript, or in the decision to publish the results.

## Notes

### Competing Interest Statement

The authors have declared no competing interest.

