## Supplemental for "Inter-Organellar Effects of Defective ER-localized Linolenic Acid Formation on Thylakoid Lipid Composition and Xanthophyll-Cycle Pigment De-epoxidation in the Arabidopsis *fad3* mutant"

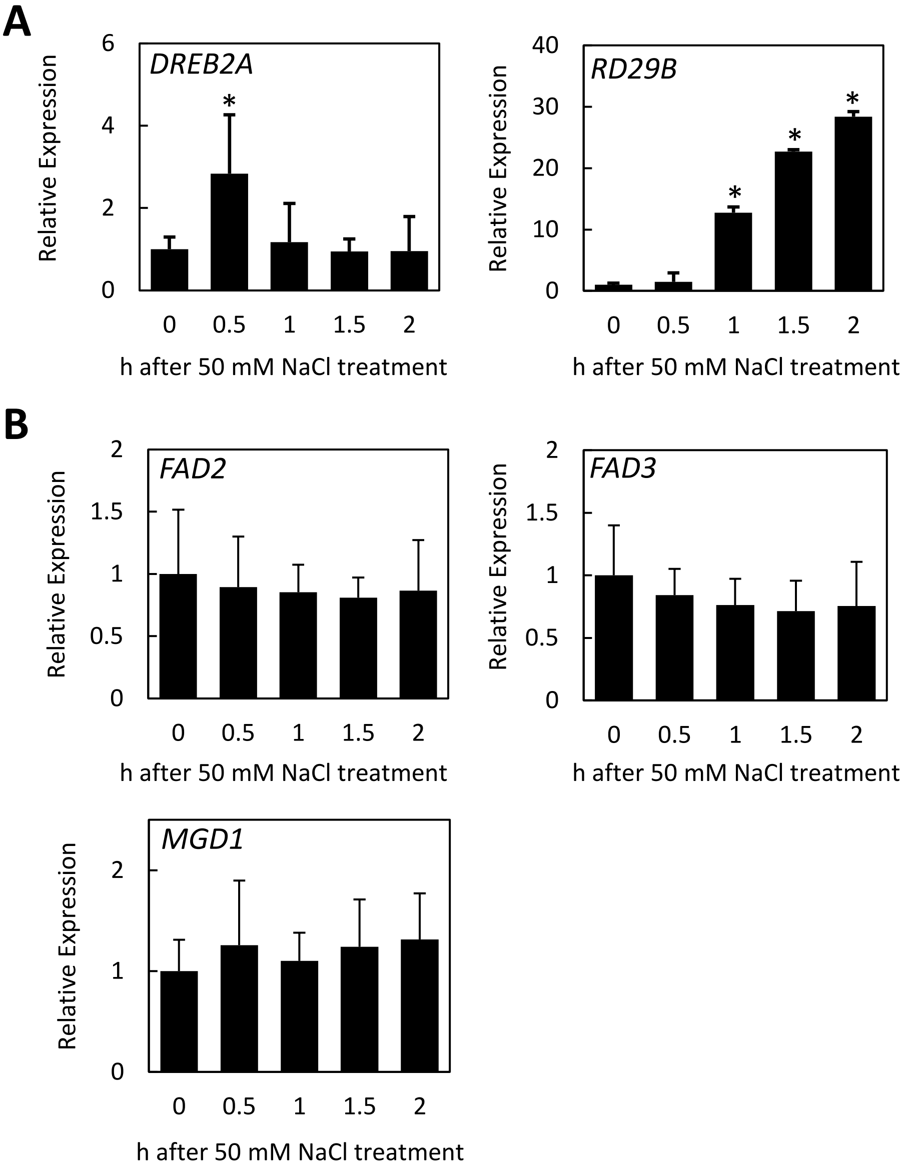


**Supplemental Fig. 1. Transcript abundance of salt stress marker genes and genes involved in lipid biosynthesis upon exposure to mild salt treatment.**

Seedlings grown on solid media for two weeks were transferred to solid media containing 50 mM and harvested after 0, 0.5, 1, 1.5 and 2 h. RNA was extracted and used for cDNA synthesis to analyze the transcript abundance of selected genes by quantitave real time PCR (qPCR). **A**, Transcript abundance over time after salt treatment for the salt-activated genes *DREB2A* and *RD29B.* **B***,* Transcript abundance over time after salt treatment for *FAD2, FAD3* and *MGD1*. Data were normalized to *UBC10* as an internal reference and represent means ± standard deviations based from two independent experiments and four biological replicates. Asterisks indicate significant changes compared to time point zero according to a Student's T-test (**P*<0.05).

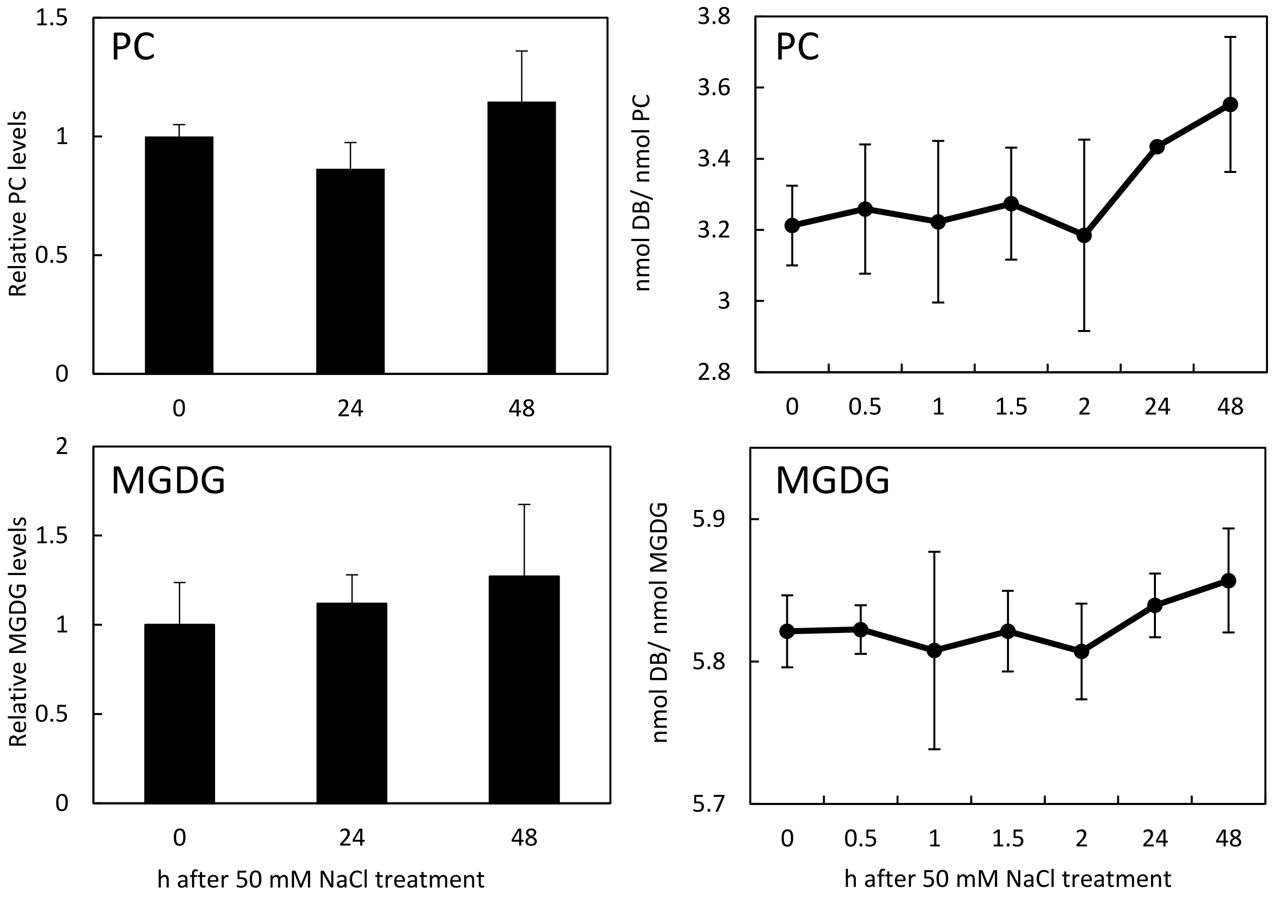


**Supplemental Figure 2. Degree of unsaturation of PC and MGDG after long-term salt treatment of wild type seedlings.**

The abundance and degree of lipid unsaturation of phosphatidylcholine (PC) and monogalactosyldiacylglycerol (MGDG) was analyzed for wild type seedlings treated for different periods with 50 mM salt. Seedlings were grown on ½ MS solid media and transferred to ½ MS solid media containing 50 mM NaCl. Green tissue was harvested after the times indicated, lipids were extracted, separated into a galactolipid and a phospholipid fraction by solid phase chromatography and then individual lipid classes were separated by thin layer chromatography. Phosphatidylcholine (PC) and monogalactosyldiacylglycerol (MGDG) were isolated and quantified according to gas-chromatographic analysis of their associated fatty acids after chemical transmethylation. Based on the fatty acid profiles of each time point displayed, the nmol double bonds (DB) were calculated. The graphs show the Analysis of wild type seedlings exposed for 24 h or 48 h of salt treatment. Left panels, Amounts of PC and MGDG, as indicated. Right panels, Degree of lipid unsaturation over 48 h of salt treatment, given as the number of double bonds per lipid for PC and MGDG, as indicated. Data represent means ± SD from three experiments performed in triplicates.

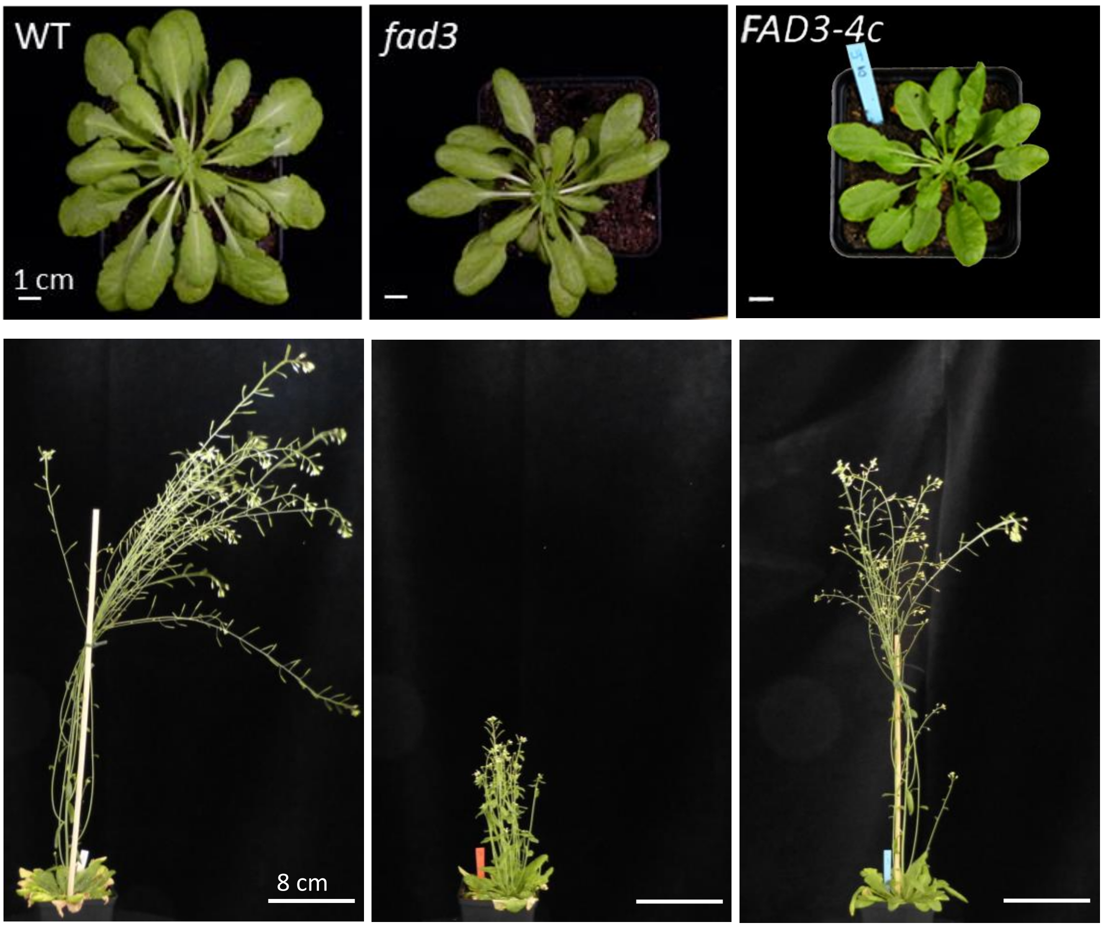


**Supplemental Figure 3. Phenotype of the *fad3* mutant under short day and low light conditions and genetic complementation.**

When grown under short day conditions (8 h light at 21 °C, 16 h dark at 18 °C) and low light intensity (170–200 µmol $m^{-2}$ $sec^{-1}$), the *fad3* mutant displayed developmental retardation. **A**, Seedlings of wild type, the *fad3* mutant and the *FAD3-4c* line were grown for two weeks on ½ MS solid media and then individually transferred to soil. Plants were documented after growth for another seven weeks. Scale, 1 cm. Documented were 4-5 individual plants. **B**, Plants grown for two weeks on ½ MS solid media were individually transferred to soil and grown for seven weeks under short-day (8 h light, 16 h dark) conditions. A difference in the formation of inflorescences was observed after transferring the plants to long-day conditions (16 h light, 8 h dark). Scale, 8 cm. Images are representative for 8-10 plants analyzed.

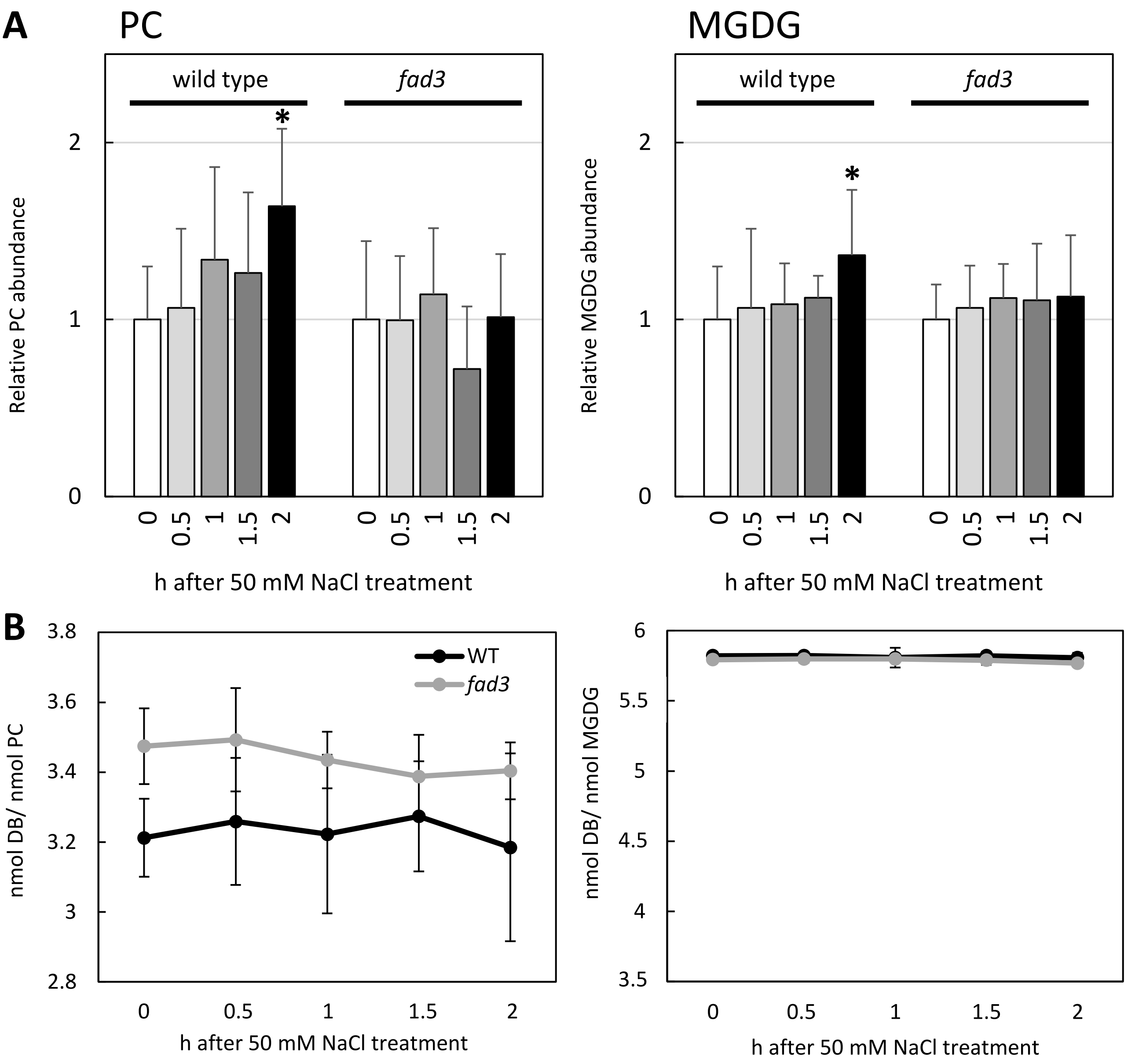


**Supplemental Figure 4. Degree of unsaturation of PC and MGDG in wild type seedlings, the *fad3* mutant and in the complemented FAD3-4c line upon exposure to mild salt treatment.**

The abundance and degree of lipid unsaturation of phosphatidylcholine (PC) and monogalactosyldiacylglycerol (MGDG) was analyzed for wild type, *fad3* or *FAD3-4c* seedlings treated for different periods with 50 mM salt. Seedlings were grown on ½ MS solid media and transferred to ½ MS solid media containing 50 mM NaCl. Green tissue was harvested after the times indicated, lipids were extracted, separated into a galactolipid and a phospholipid fraction by solid phase chromatography and then individual lipid classes were separated by thin layer chromatography. Phosphatidylcholine (PC) and monogalactosyldiacylglycerol (MGDG) were isolated and quantified according to gas-chromatographic analysis of their associated fatty acids after chemical transmethylation. Based on the fatty acid profiles of each time point displayed, the nmol double bonds (DB) were calculated. **A**, Seedlings of wild type controls and *fad3* mutants were subjected to 50 mM salt treatment and the relative increases of PC (left panel) and MGDG (right panel) were analyzed after the times indicated. The horizontal line indicates the normalized background (set as 1). B, The degree of lipid unsaturation was calculated from data shown in (A) is given as the number of double bonds per lipid for PC (left panel) and MGDG (right panel), as indicated. Data represent means ± SD from 3 experiments with 1-6 biological replicates. Asterisks indicate a significant change compared to time point 0 of the respective plant line, according to a Student's T-test (*P<0.05).
